# RNA sequencing analyses of gene expression by CRISPR/Cas9 knockout of CLL-1 gene in acute myeloid leukemia cells

**DOI:** 10.1101/2022.11.10.515971

**Authors:** Yanyu Wang, Songfang Wu, Jing Yang, ShiLiang Wang, Hong Xiong, Shui-Jun Li

## Abstract

CLL-1 has been revealed its potential role in acute myeloid leukemia (AML), however, the underlying mechanisms remain unclear. CRISPR/Cas9 strategy was employed to knock out CLL-1 gene in U937 cells and western-blot was used to validate the success of knock out. CCK8 and Transwell assays were used to detect cells viability and migration, respectively. RNA-sequencing was performed to profile mRNA expression in CLL-1 gene knock-out and wide type U937 cells. A cutoff of 1.5-fold change and false discovery rate (FDR) <0.05 was used to screen differentially expressed genes (DEGs), which were presented by volcano plots and hierarchical cluster heatmap. Protein-protein interaction (PPI) network was constructed by String database and Cytoscape software. Furthermore, hub genes were mined by CytoNCA and MCODE, which were subjected to functional enrichment using R package. Finally, the findings were validated using qRT-PCR and western-blot. The protein level of CLL-1 was significantly lowered, and cell viability and migration were suppressed in knock-out cells compared to wide type. Using RNA-sequencing and bioinformatics analysis, 452 DEGs (179 up-regulated and 273 down-regulated) were obtained, and several important hub genes (such as CCR2, FBXO21, UBB and UBE2C) were filtered out, which were enriched in 132 GO terms and 36 KEGG pathways such as chemokine signaling pathway and ubiquitin mediated proteolysis. A total of 8 representative genes mRNA expressions were validated by qRT-PCR, and the protein levels of 6 genes were confirmed by western-blot. CLL-1 gene might exert its role in AML through modulating genes enriched in multiple functions such as chemokine signaling and ubiquitination. Our results may give us new knowledge of CLL-1 in AML and provide a basis for mining novel targets.

## Introduction

Acute myeloid leukemia (AML) is a disease of the bone marrow, a disorder of hematopoietic stem cells due to genetic alterations in blood cell precursors resulting in overproduction of neoplastic clonal myeloid stem cells (Pelcovits & Niroula, 2020). AML is the most common acute leukemia in adults and among the most lethal (Kadia, Ravandi, Cortes, & Kantarjian, 2016). The median age at diagnosis of AML is around 70 years, and approximately 3% of AML cases occur in children age 14 years or younger (Tamamyan et al., 2017). The 2018 AML incidence estimates from SEER are <1.23 per 100,000 in the <40-year-old population, 10.92 per 100,000 in the ≥60-year-old population and 20.89 per 100,000 in the ≥75-year-old population in the USA (Lin, Zhang, Yu, & Wu, 2021). However, there has been little progress in the standard therapy for AML over the past four decades (Yang & Wang, 2018), further understanding of the molecular heterogeneity and pathogenesis of AML is needed to develop novel therapies.

With unprecedented advances in molecular genetics, a deeper insight into the biology has opened doors to the development of therapeutic approaches in the hope of achieving durable remissions and improving survival (Higgins & Shah, 2020). Besides, recent advances in immunotherapy have generated substantial excitement for cancer patients. Due to the immunosuppressive nature of AML, the activation of the immune system through genetically engineered T-cell therapy presents a promising, curative option for patients (Gill, 2019). C-type lectin-like receptors play a pivotal role in the fight against infection and maintain homeostasis and self-tolerance by recognizing damage associated and pathogen associated molecular patterns leading to regulation of innate and adaptive immunity (Ma, Padmanabhan, Parmar, & Gong, 2019).C-type lectin-like molecule-1(CLL-1 or CLEC12A) is a type-II transmembrane glycoprotein, which belongs to the C-type lectin-like receptor family (J. Wang et al., 2018). It is reported that CLEC12A/CLL-1 played an essential role in attenuating sterile inflammation which is induced by uric acid crystal in a Syk-dependent pathway (Neumann et al., 2014). In a collagen antibody-induced arthritis (CAIA) model, Clec12a^-/-^ mice experienced more severe inflammation during CAIA due to the over-activation of myeloid cells (Begun et al., 2015).

Intriguingly, CLEC12A has been found selectively present on leukemic stem cells (LSCs) in AML but absent in normal HSCs (Leipold et al., 2018), which may be an effective alternative target for AML with specificity against leukemic progenitor cells and their progeny, while sparing normal myeloid precursor cells. Indeed, mono-antibody therapy targeting CLL-1 has been revealed its potential efficacy against AML cells and shown to be effective in reducing AML burden in xenograft model (Lu et al., 2014). Researchers have developed and optimized CLL-1 CAR-T for AML and showed efficient and specific anti-leukemia activity to AML cell lines and primary blasts from AML patients, as well as in mouse model (Laborda et al., 2017; Tashiro et al., 2017). Our previous study revealed that CLL-1 is a novel prognostic predictor that could be exploited to supplement the current AML prognostic risk stratification system, and potentially optimize the clinical management of AML (Y. Y. Wang et al., 2017).However, the exact physiological function and underlying mechanisms of CLL-1 in AML need to be elucidated.

In the present study, CRISPR/Cas9 was used to knock out CLL-1 gene in AML U937 cells, and high-throughput RNA sequencing was performed to profile the changes of mRNA expression. The results will broaden our knowledge of CLL-1 gene’s function and provide novel therapeutic targets in AML.

## Methods

### Cell culture

The AML cell line U937 was obtained from the American Type Culture Collection (Rockville, MD, USA), and maintained at 37 °C, in the presence of 5% CO2 and in a humidified atmosphere. The cells were cultured in RPMI-1640 medium (Gibco Technologies, Germany), supplemented with 10% fetal bovine serum (Moregate, Australia) and 1% penicillin/streptomycin (10,000 U/ml and 10,000 μg/ml respectively; Gibco/Life Technologies, Germany).

### Knockout of CLL-1 gene using CRISPR/Cas9

CLL-1 gene was knocked out by Bioray Laboratories Inc. (Shanghai, China) using the CRISPR/Cas9 gene editing system according to manufacturer’s instructions. In brief, the primers including CLL-1 sgRNA1-forward (5’-CACCGGCTGGACGC CATACATG AGA-3’), CLL-1 sgRNA1-reverse (5’-AAACTCTCATGTATGGCGT CCAGC-3’), CLL-1 sgRNA2-forward (5’-CACCGGATATAGCTCACGACATAATT-3’) and CLL-1 sgRNA2-reverse (5’-AAACAATTATGTCGTGAGCTATATC-3’) were synthesized. Cloning of the sgRNA oligos was performed using lentiCRISPR V2 vector. The vector lentiCRISPR V2 was digested using BbsI restriction endonuclease. The diluted sgRNAs were then ligated in the vector lentiCRISPR V2 with T4 DNA ligase. The ligated vector was inserted into E.cloni® 10G electro-competent cells. Plasmid construction was performed according to protocol and confirmed by sequencing. The U937 cells were transfected with CLL-1 knock-out plasmid using Lipofectamine 2000 for 48 h. The transfected cells were selected with puromycin (3ug/ml) for seven days. Thereafter, single-cell cloning was performed in a 96-well plate to grow single clones. After growth, western blot and sequencing were performed to confirm knock-out of CCL-gene according to manufacturer’s protocols.

### Cell viability assays

The CCK-8 assay (Dojindo Molecular Technologies, Gaithersburg, MD, USA) was performed to evaluate cell viability according to manufacturer’s instructions. Briefly, cells were seeded into 96-well plates (5×10^4^ cells/well) and cultured for 24h,48 h and 72h, respectively. Subsequently, 10 μl of CCK-8 solution was added to each well and incubated at 37 °C for 1 h. The absorbances (Abs) at 450 nm were recorded using a microplate reader (Bio-Rad, Hercules, CA, USA).

### Cell migration assay

Cell migration was assessed using Transwell chambers with 8 μm pore size membrane inserts (BD Falcon™; BD Biosciences) according to previously described methods (Han et al., 2018). Briefly, 2×10^5^ cells were seeded into the upper chamber supplemented with 100 μl serum-free medium, while 700 μL RPMI1640 containing 10% FBS was added to the lower chamber. After 12 hours’ incubation (5% CO_2_, 37°C), non-migrated cells in the upper chamber were removed and migrated cells in the lower chamber were fixed with 4% paraformaldehyde and stained with 0.1% crystal violet. Finally, cells were washed with PBS and counted using an inverted microscope (Olympus Corporation) with ImageJ software. The experiments for each group were repeated in triplicate.

### RNA sequencing raw data acquisition

CLL-1 gene knock-out (KO) and wide type (WT) U937 cells (n=3, each group) were harvested and subjected to total RNA extraction using TRIzol reagent (Beyotime, China). RNA quality was evaluated by an Agilent 2100 Bioanalyzer, and RNA samples (n=3, each group) with RNA integrity number (RIN) more than 7 were used for purification, library preparation, amplification and sequencing for 150bp paired end reads using Illumina HiSeq 2500 platform according to manufacturer’s instructions.

### RNA sequencing data analysis

Before read mapping, clean reads were obtained from the raw reads by removing the adaptor sequences and low-quality reads using FastQC software. The clean reads were then aligned to Human genome (GRCh38, NCBI) using the Hisat2 (Kim, Langmead, & Salzberg, 2015) and reconstructed by Cufflinks (Ghosh & Chan, 2016). HTseq was used to calculate the expression of genes (Anders, Pyl, & Huber, 2015). The read counts of each transcript were normalized to the length of the individual transcript and to the total mapped read counts in each sample and expressed as FPKM (Fragments Per Kilobase of exon per Million mapped reads). Differential expression analysis was performed using DESeq2 with the threshold of 1.5-fold change (| log_2_fold change | >0.58) and false discovery rate (FDR) less than 0.05. The differential expressed mRNAs between WT and KO U937 cells were presented with volcano plots and hierarchical cluster heatmap using R software.

### GO and KEGG pathway enrichment

To reveal the function of the differential expressed genes between WT and KO U937 cells, gene ontology (GO) (including biological process, cellular component and molecular function) and Kyoto Encyclopedia of Genes and Genomes (KEGG) pathway analysis was performed using the Bioconductor package clusterProfiler.

### Protein-protein interaction network and module analysis

Protein-protein interaction (PPI) was screened using STRING (https://string-db.org/) database with interaction score of 0.9 as the threshold, and the PPI network was visualized by Cytoscape software. Furthermore, CytoNCA was used to screen hub genes based on the nodes degree, and the candidate modules were obtained by molecular complex detection (MCODE) with default parameters: degree cut-off = 2, node score cut-off = 0.2, *k*-core = 2, and max depth = 100.

### Real-Time qPCR

Total RNA was extracted from WT and KO U937 cells using TRIzol reagent (Beyotime, China). Real-time qPCR was performed according to previously described methods (Zhu et al., 2019). Briefly, the cDNA was synthesized using the PrimeScript RT reagent Kit (Yeasen, China) and amplified by real-time qPCR with an SYBR Green Kit (Yeasen, China) on QuantStudio 12K Flex Real-Time PCR System. The relative gene expression levels were determined using 2^-ΔΔCt^ method with GAPDH as an internal control. All of the primers were synthesized by Biosune (Shanghai, China), and the sequences of primers were shown in Table 1.

### Western-blot

WT and KO U937 cells were harvested and subjected to total protein extraction using RIPA lysis. For each sample, 50 μg of protein was used for gel electrophoresis in 10-14% SDS-PAGE gels and transferred to PVDF membranes (Merck Millipore, Billerica, MA, USA). After blocking in 5% defatted milk, the membranes were incubated with primary antibodies of FBXO21, FBXO21, FBXO25, UBB, USP53, UBE2C, UBE2E3, USP2, USP44 and B-Actin overnight at 4 °C. After incubation with secondary antibodies, signals were detected using the ECL detection system (Thermo Fisher Scientific) and analyzed by ImageJ software.

### Statistical analysis

The data are presented as the mean ± SD, and statistical analysis was performed using SPSS software (version 22.0). Student’s t-test was used to compare continuous variables, and P < 0.05 were considered as statistically significant.

## Results

### Confirmation of CLL-1 gene knock-out

CLL-1 gene knock-out was confirmed by western blotting (GAPDH serve as internal control) to ensure the absence of CLL-1 protein expression. As shown in Figure 1, the CLL-1 protein level was significantly lowered in CLL-1 gene knock out U937 cells compared to wide type, which indicated the successful knock-out of CLL-1 gene.

**Figure 1.**
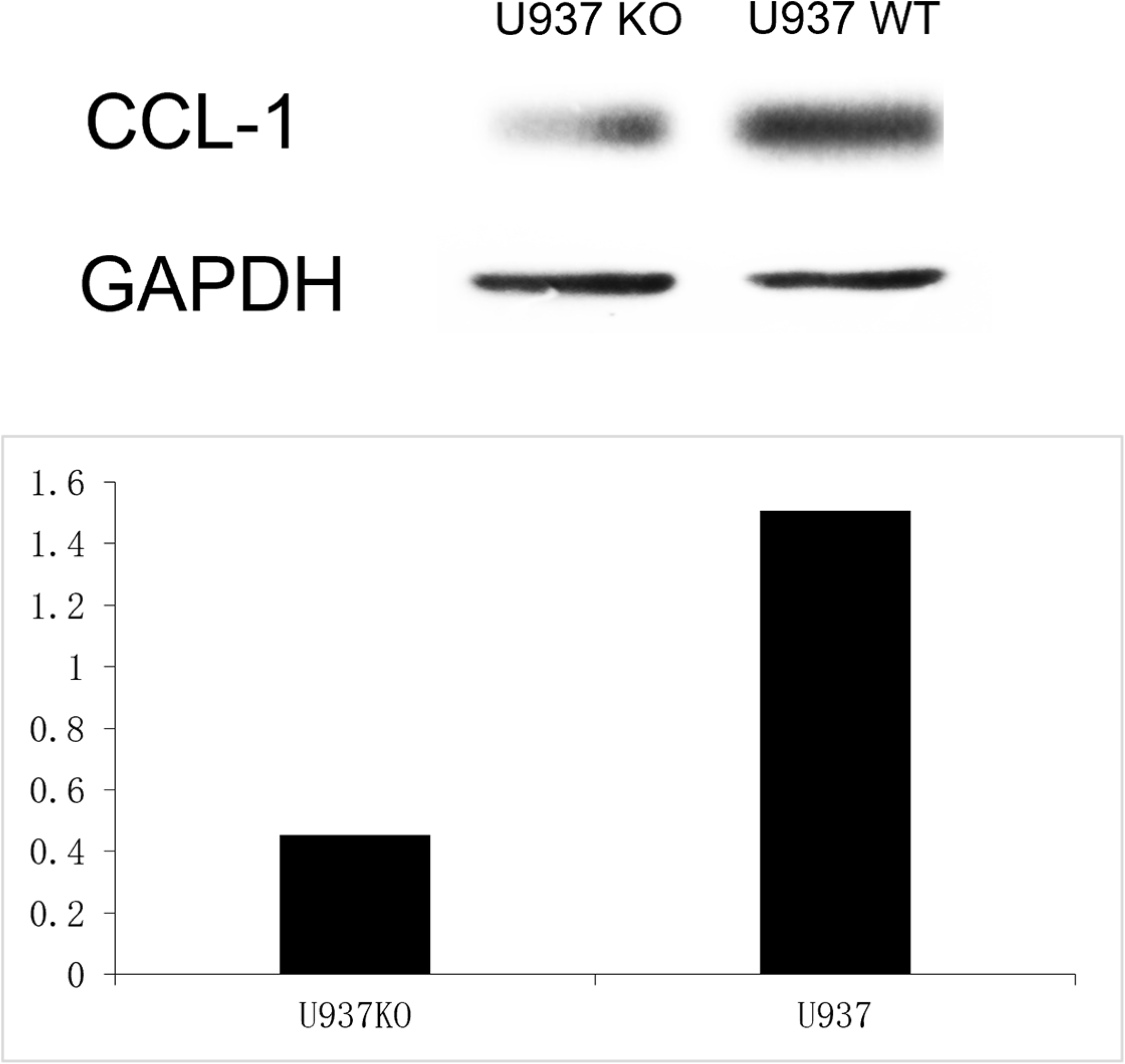
Western-blot analysis of CLL-1 expression in knock-out and wide type U937 cells.

### CLL-1 gene knock-out reduced cells viability and migration

To examine the effects of CLL-1 gene knock-out on U937 cells viability and migration, CCK-8 and Transwell assays were performed, respectively. After CLL-1 gene knock-out, the cells viability was obviously reduced at the time-point of 24h,48h and 72h(Figure 2A). Besides, the migrated cells in CLL-1 knock-out group were significantly decreased compared to wide type group (Figure 2B). These results indicated CLL-1 gene might play an important role in the cell viability and migration of AML.

**Figure 2.**
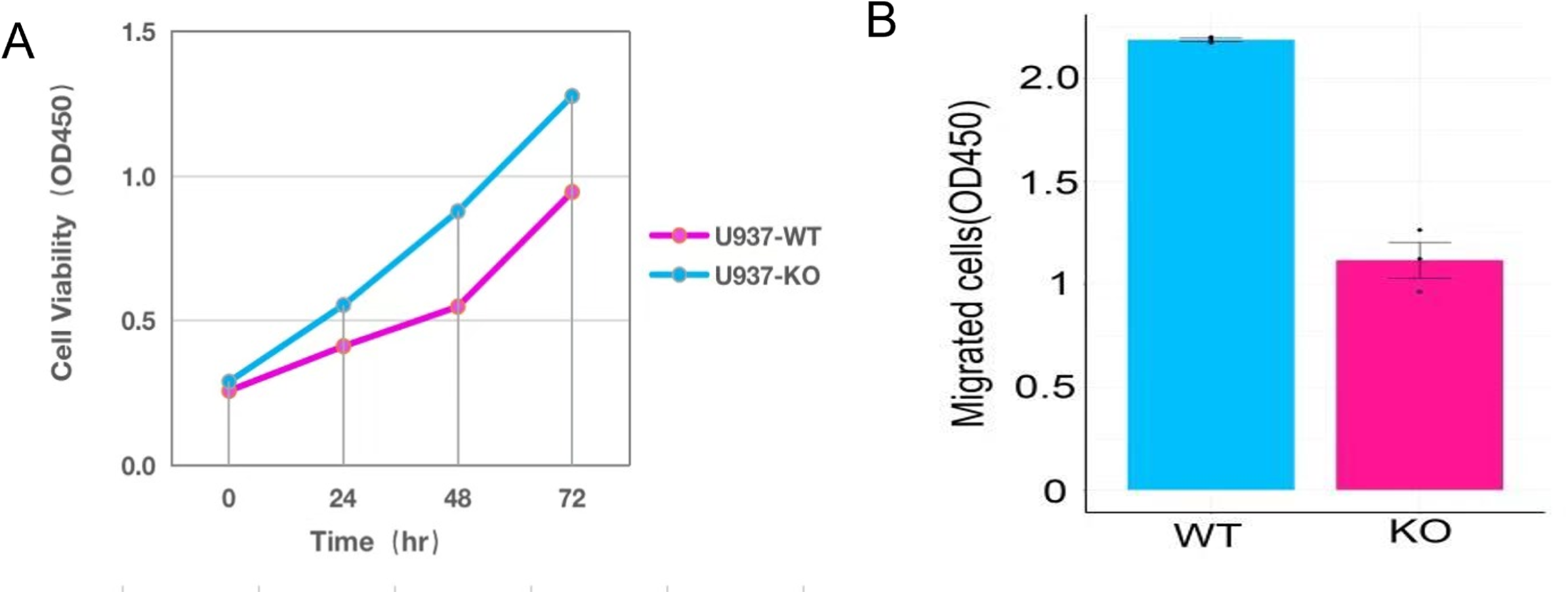
CCK8 and Transwell assays in CLL-1 knock-out and wide type U937 cells. A. CCK8 assay, B. Transwell assay.

### CLL-1 gene knock-out altered mRNA profile in U937 cells

To explore the underlying mechanisms of CLL-1 affecting AML, RNA-seq was performed to profile the mRNA expression in CLL-1 knock-out and wide type U937 cells. The differential gene analysis (Figure3 A & B) revealed that there were 452 differentially expressed genes (DEGs) in KO cells compared to WT cells. In addition, we noticed that there were 179 up-regulated (including TRMT12, IL1B, CD86, etc.) and 273 down-regulated mRNAs (including FBXO25, FBXO21, UBE2C, etc.) The top 30 representative mRNAs were listed in Table 2, and 452 DEGs were listed in Table S1.

**Figure 3.**
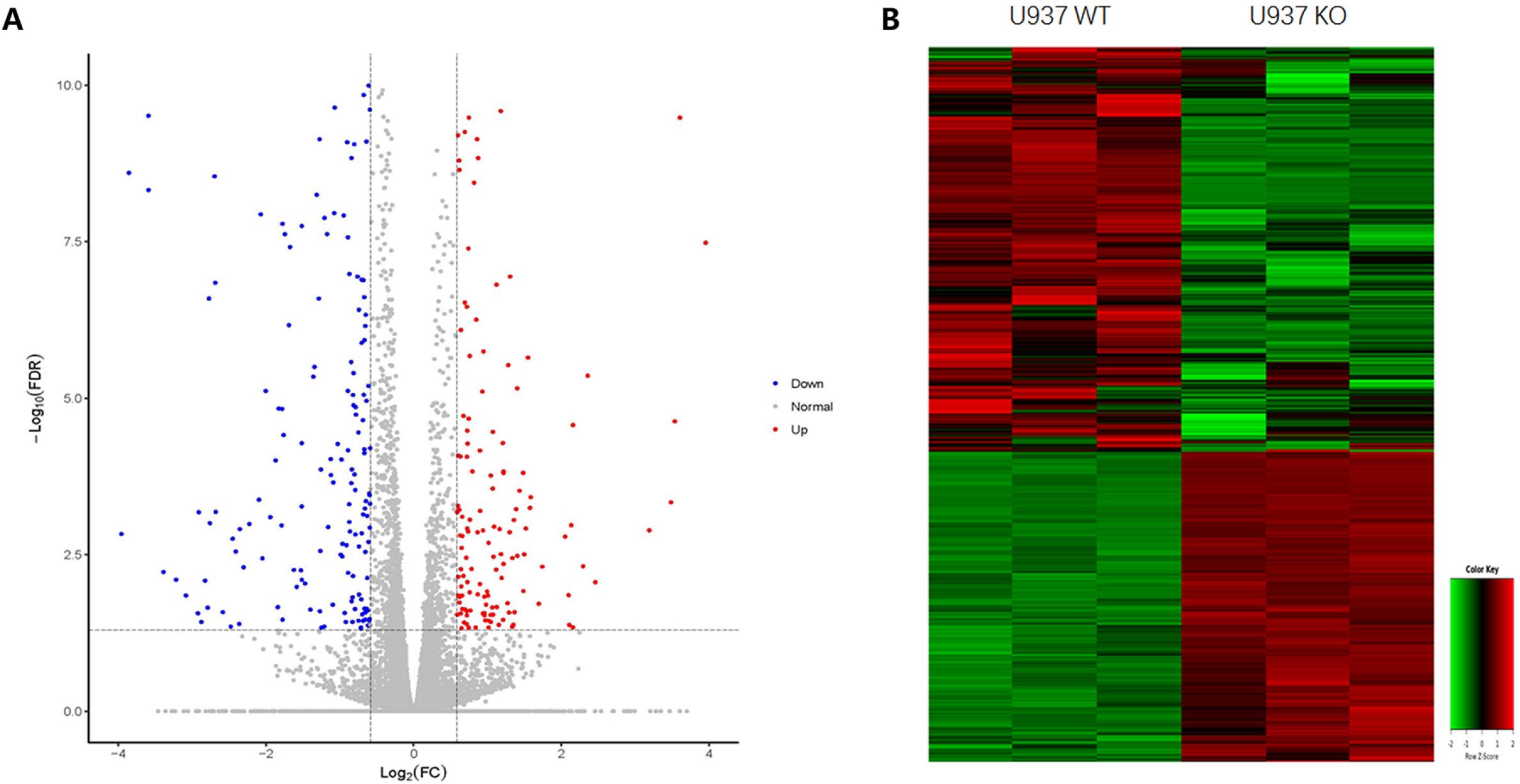
Volcano plots and hierarchical cluster analysis. A. Volcano plots, blue dots represent down-regulated mRNAs, grey dots represent no significant changed mRNAs, red dots represent up-regulated mRANs. B. Heatmap of hierarchical cluster, red colors indicate up-regulated and green colors indicate down-regulated.

### Protein-protein interaction network analysis

To reveal protein-protein interaction (PPI) among DEGs, 452 DEGs were imported to String database to screen and visualized by Cytoscape software. A PPI network with 340 nodes and 904 edges was obtained (Figure 4). To obtain the hub genes in the PPI network, CytoNCA was performed. As shown in Figure 5A, the top 30 genes with higher degrees (>13) were filtered out, which included UBB, IL1B, CD86, etc. Furthermore, MCODE was performed to explore the important modules in the PPI network. As shown in Figure 5B, we obtained a module with the highest socre (9.684), which composed of 20 nodes (including FBXO21, UBB, UBE2C, etc.) and 92 edges.

**Figure 4.**
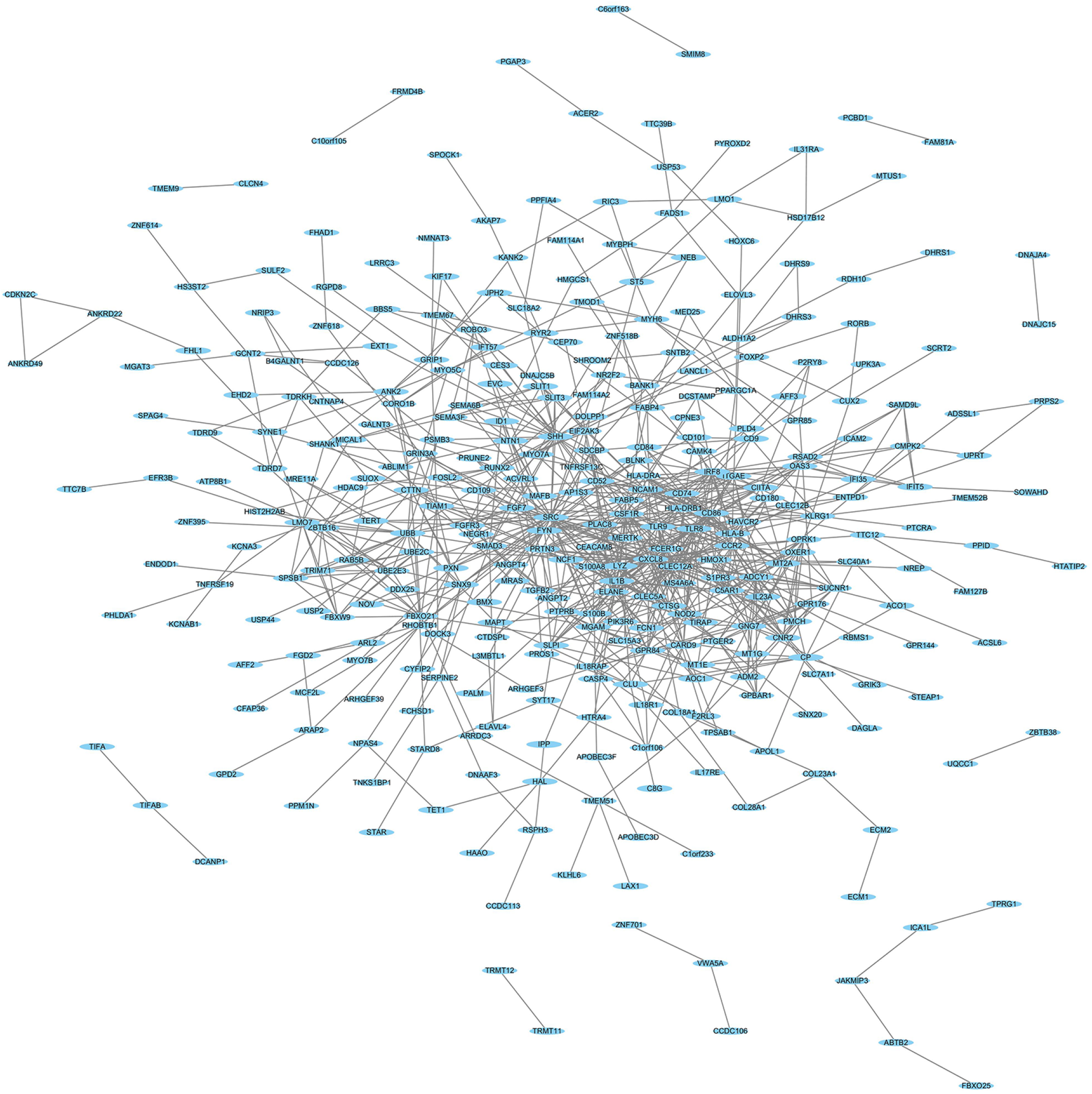
PPI network of 452 DEGs. PPI network with 340 nodes and 904 edges, and blue nodes indicate DEGs and edges indicate interactions.

**Figure 5.**
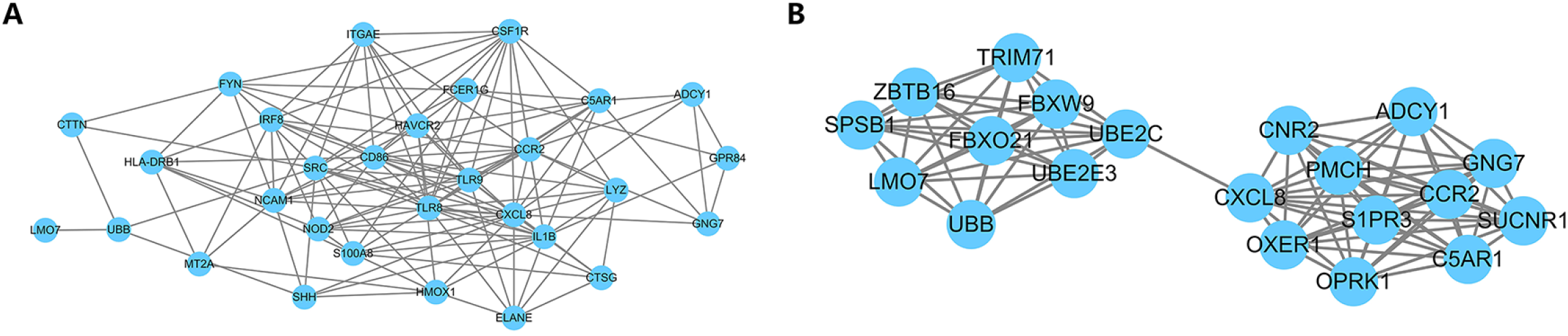
Subnetworks of hub genes and an important module. A. Subnetwork of the top 30 hub genes with higher degrees (>13). B. Subnetwork of an important module with the highest socre (9.684). Blue nodes indicate DEGs and edges indicate interactions.

### GO and KEGG pathway functional enrichment

To better understand the biological functions of the 20 DEGs involved in the important module, functional enrichment was performed. We noticed that these DEGs were enriched in 132 GO terms and 36 pathways, including ubiquitin mediated proteolysis, chemokine signaling pathway, protein ubiquitination, etc. For example, GNG7, ADCY1, CXCL8 and CCR2 were enriched in chemokine signaling pathway. UBE2C and UBE2E3 were enriched in ubiquitin mediated proteolysis and genes including SPSB1, ZBTB16, UBB, TRIM71, UBE2C, FBXO21, LMO7 and UBE2E3 were enriched in protein ubiquitination. The TOP 30 GO terms and KEGG pathways were presented in Figure 6, and detailed information was listed in Table S2 and Table S3.

**Figure 6.**
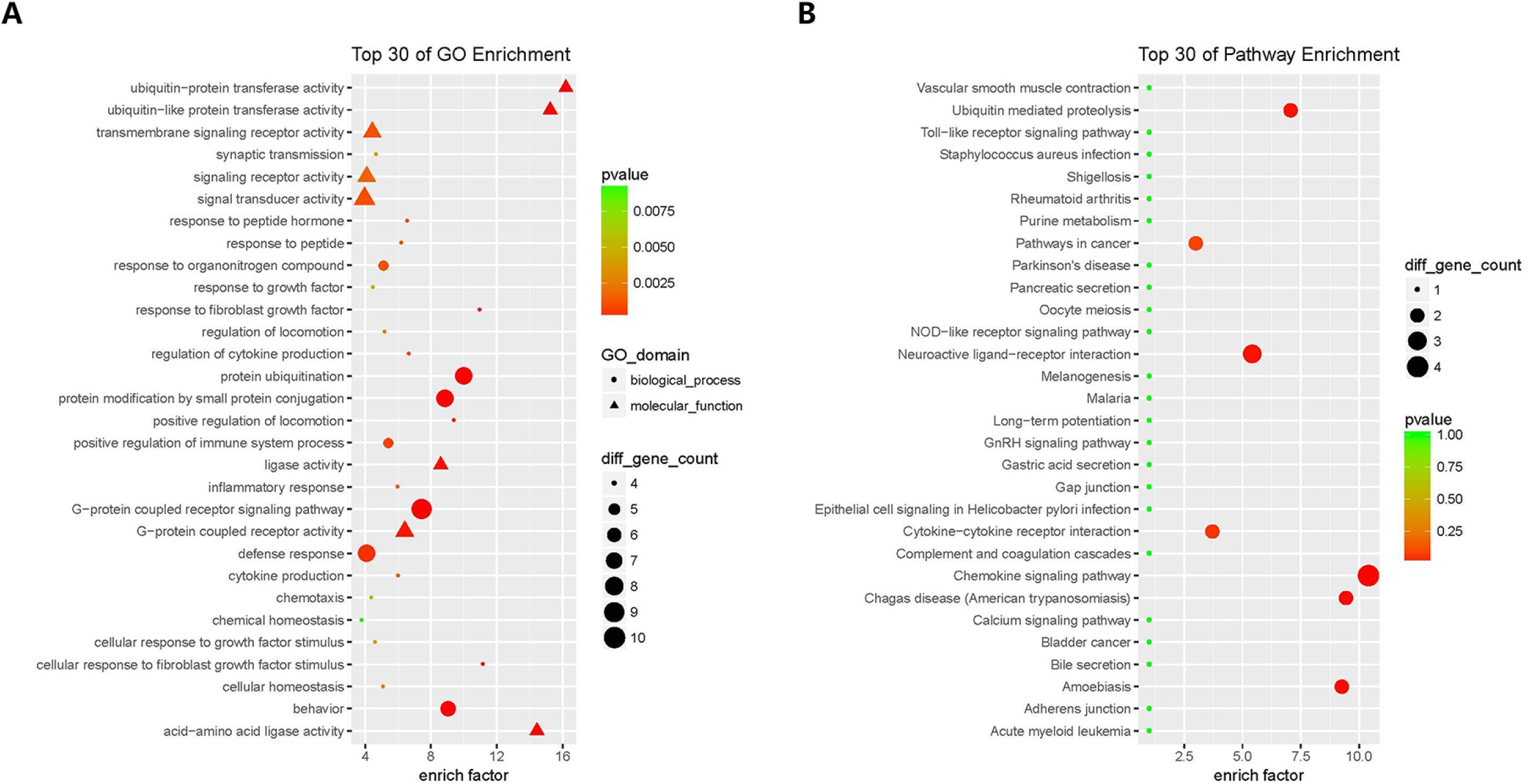
Top 30 GO terms and KEGG pathways. A. Top 30 GO terms, B. Top 30 KEGG pathways.

### Validation of RNA-sequence data by qRT-PCR and western-blot

To validate our findings from RNA-sequence, a total of 8 genes (including FBXO21, FBXO25, UBB, USP53, UBE2C, UBE2E3, USP2 and USP44) were selected to perform qRT-PCR and western-blot experiments. The results of qRT-PCR and RNA-sequence data of all the 8 genes were in good accordance (Figure 7A), and 6 genes (FBXO21, FBXO25, UBB, USP53, UBE2C and UBE2E3) expression were further verified in the protein level. These results indicated the reliability of our RNA-sequence analysis.

**Figure 7.**
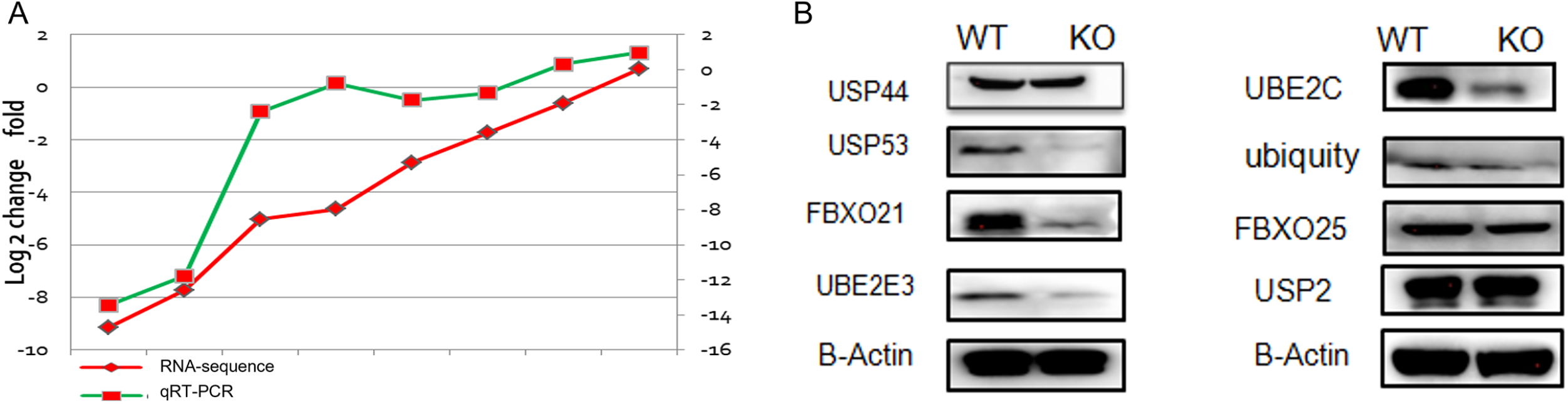
Validation by qRT-PCR and western-blot. A. The mRNA levels of 8 genes were validated by qRT-PCR, the red line represents RNA sequence data and green line represents qRT-PCR data. B. The protein levels of 8 genes were validated by western-blot.

## Discussion

In the present study, we aimed to explore the effects and underlying mechanisms of CLL-1 gene in AML, RNA-sequencing was performed to profile mRNA expression in CLL-1 gene knock-out and wide type U937 cells. We obtained numerous differentially expressed mRNAs in CLL-1 gene knock-out cells, and filtered out a batch of important genes which enriched in multiple GO terms and KEGG pathways.

It is reported that CLL-1 gene might be the most prominently differently expressed surface markers in AML (Daga et al., 2019). We observed that the viability and migration were reduced in CLL-1 gene knock-out U937 cells, which implicated the important role of CLL-1 in AML. Using RNA-sequencing, we obtained a batch of differentially expressed genes related to CLL-1 gene knock-out, which might account for the underlying mechanisms of its role in AML. AML is a bone marrow disease in which the leukemic cells show constitutive release of a wide range of CCL and CXCL chemokines and express several chemokine receptors (Kittang, Hatfield, Sand, Reikvam, & Bruserud, 2010). It is noticed that chemokine receptors may play an important role in orchestrating the migration of cells and mediating the recruitment of immune cells (Stone, Hayward, Huang, Z, & Sanchez, 2017). CC chemokine receptor 2 (CCR2) is the chemokine receptor, which has been found to be associated with advanced cancer, metastasis, and relapse (Nagarsheth, Wicha, & Zou, 2017). It is reported that tumor growth was reduced in CCR2 ^-/-^ mice compared to wild-type mice (Huang et al., 2007). It was shown that CCR2 was almost exclusively expressed on monocytoid AML in human samples (Cignetti et al., 2003). High expression of CCR2 was observed in AML cell lines and 65% of human AML samples, and the blockade of CCL2/CCR2 axis inhibited cells transmigration and proliferation in AML (Macanas-Pirard et al., 2017). These previous studies indicated that chemokine receptor CCR2 might play an important role in AML. In the present study, we noticed that CCR2 gene expression was significantly lowered in CLL-1 knock-out U937 cells, which was partially consistent with previous studies and implicated that CLL-1 gene might affect AML through CCR2 signaling. However, we did not observe the change of CCR2 ligand, which needed further investigation. In addition, we also observed several hub genes (such as GNG7, ADCY1and CXCL8) enriched in chemokine signaling pathway were altered, which suggested that CLL-1 gene might modulate chemokine signaling in AML.

Besides, we noticed that a batch of important genes such as UBB, UBE2C, FBXO21, USP53 and UBE2E3 were significantly changed in CLL-1 knock-out cells and enriched in ubiquitination related GO terms and pathways, which indicated that CLL-1 might regulate ubiquitination in AML. Ubiquitination is an essential post-translational modification involved in protein stability, localization, interactions, and activity, which influences cell apoptosis, cell survival, cell-cycle progression, DNA repair and antigen presentation, and is implicated in multiple pathophysiological states and diseases such as cancer, infections and hereditary disorders (van Wijk, Fulda, Dikic, & Heilemann, 2019).Ubiquitination is mediated by the sequential action of ubiquitin-activating enzyme, ubiquitin-conjugating enzyme and ubiquitin protein ligase (Manasanch & Orlowski, 2017), and the abundance of cellular ubiquitin is modulated by a family of multiple ubiquitin genes such as UBB and UBC (Haakonsen & Rape, 2017). UBB has been implicated in many tumors including ovarian cancer, gastric cancer, lung adenocarcinoma, etc (Deng, Huang, Wang, & Chen, 2020; Gong, Lin, & Yuan, 2020; Scarpa et al., 2020). However, there is no report about UBB in AML. Here, we reported that the expression of UBB mRNA and protein levels were lowered in CLL-1 knock-out U937 cells, which indicated that UBB might play a role in AML. It has been reported that UBE2C (an ubiquitin-conjugating enzyme) participated in carcinogenesis by regulating the cell proliferation, apoptosis, and transcriptional processes (Jin et al., 2019). UBE2C mRNA and/or protein levels were aberrantly increased in many cancer types with poor clinical outcomes, and inhibition of UBE2C suppressed proliferation, clone formation, and malignant transformation in tumor cells (Xie, Powell, Yao, Wu, & Dong, 2014). Although little is known about UBE2C in AML, it has been reported that UBE2C gene expression was increased in aneuploid acute myeloid leukemia, which implicated the correlation of UBE2C with AML (Simonetti et al., 2019). In addition, FBXO21 belongs to F-box proteins which serves as the substrate-recognition subunit of a SKP1-CUL1-F-box protein (SCF)-type ubiquitin ligase (Watanabe, Yumimoto, & Nakayama, 2015), and ubiquitin-specific proteases 53 (USP53) belongs to the family of deubiquitinating enzymes, which catalyze the reversible modification of target proteins with ubiquitin and stabilize proteins (Fraile, Quesada, Rodriguez, Freije, & Lopez-Otin, 2012). It has been reported that knockdown of USP53 in Siha cells downregulated damage-specific DNA binding protein and caused G2/M cell cycle arrest and decreased the survival rate of cells in response to radiation (Zhou, Yao, Wu, Chen, & Fan, 2020). However, little has been reported about the role of FBXO21 and USP53 in AML. In the present study, we noticed that the mRNA and protein levels of FBXO21 and USP53 were significantly decreased after CLL-1 gene knock-out. Further investigations on these genes enriched in ubiquitination related GO terms and pathways may provide novel targets in AML.

Despite the significant findings in the present study, there are some limitations should be noted. Although several important genes expression were validated in mRNA and protein levels, our results were mainly based on RNA-sequencing and bioinformatics analysis, which need further biological function investigation. Besides, our results were obtained from AML cell models, which needed further animal models and large-scale patient experiments to verify.

In conclusion, knock out of CLL-1 gene suppressed cell viability and migration in AML cell line U937 cells, and the underlying mechanisms might be related to a batch of important genes which were enriched in multiple functions such as chemokine signaling and ubiquitination. Our results may give us new knowledge of CLL-1 in AML and provide a basis for mining novel targets.

## Acknowledgements

Not applicable.

## Supporting Information

Table S1: WSF-KO_vs_WSF-WT.log2FC0.585.FDR0.05.Diff.mRNA&circRNA

Table S2: Negative_Analysis_miRNA_mRNA.GO-Analysis

Table S3: Pathyway Analysis Result

